# Integrin signaling *via* actin cytoskeleton activates MRTF/SRF to entrain circadian clock

**DOI:** 10.1101/2021.08.12.456061

**Authors:** Xuekai Xiong, Weini Li, Jin Nam, Ke Ma

**Affiliations:** Department of Diabetes Complications & Metabolism, Beckman Research Institute of City of Hope, Duarte, CA 91010; Department of Bioengineering, University of California at Riverside, Riverside, CA 92521

**Keywords:** Circadian clock, Actin cytoskeleton, Serum Response Factor, Integrin, Myocardin-related Transcription Factor

## Abstract

The circadian clock is entrained to daily environmental cues. Integrin-linked intracellular signaling *via* actin cytoskeleton dynamics transduces cellular niche signals to induce Myocardin-related Transcription Factor (MRTF)/Serum Response Factor (SRF)-mediated transcription. So far, how the integrin-associated signaling cascade may transmit cellular physical cues to entrain circadian clock remains to be defined. Using combined pharmacological and genetic approaches, here we show that the transcription factors mediating integrin to actin cytoskeleton signaling, MRTF-A and SRF, exert direct transcriptional control of core clock components, and that this signaling cascade modulates key properties of clock circadian activity. Pharmacological inhibition of MRTF/SRF activity by disrupting actin polymerization significantly augmented clock amplitude with period shortening, whereas an actin polymerizing compound attenuated oscillation amplitude with period lengthening. Genetic loss-of-function of *Srf* or *Mrtf* mimics that of actin-depolymerizing agents, validating the role of actin dynamics in driving clock function. Furthermore, integrin-mediated focal adhesion with extracellular matrix and its downstream signaling modulates the circadian clock, as blockade of integrin, focal adhesion kinase or Rho-associated kinase (ROCK) increased clock amplitude and shortened period length. Mechanistically, we identify specific core clock transcription regulators, *Per1, Per2* and *Nr1d1*, as direct target genes of MRTF-A/SRF. Collectively, our findings uncovered an integrin-actin cytoskeleton-MRTF/SRF signaling cascade in linking clock entrainment to its extracellular microenvironment, which may mediate cellular adaptation to its physical niche.

**Author Summary:** The circadian clock anticipates and adapts to environmental changes. Interestingly, serum, as a universal clock synchronizing signal, drives intracellular actin cytoskeleton reorganization through modulation of MRTF/SRF activity. However, mechanisms that may transduce extracellular niche signals to circadian clock remains to be defined. We hypothesize that integrin-mediated intracellular signaling to actin cytoskeleton links extracellular microenvironment with MRTF/SRF transcriptional regulation to control clock function. Using small molecules and genetic approaches targeting distinct steps of integrin-actin cytoskeleton-MRTF/SRF signaling cascade, we uncover the effects of this pathway in controlling circadian clock oscillation. We also identify specific core clock regulators as direct gene target genes of MRTF and SRF-mediated transcriptional control. Our study revealed how integrin-mediated cellular interaction with its physical environment influences its intrinsic clock properties through signaling transduction *via* actin cytoskeleton remodeling, and that this mechanism may facilitate circadian clock adaptation to cellular physical niche.

## Introduction

The circadian clock has evolved as a time-keeping mechanism to anticipate and adapt to daily environmental changes essential for organismal fitness and survival (1, 2). Under physiological conditions, a central clock pacemaker residing in suprachiasmatic nuclei (SCN) of the hypothalamus is entrained by daily light input that synchronizes clock circuits within peripheral tissues. In additional to SCN-driven signals for synchronization, cell-autonomous peripheral oscillators in tissues outside SCN respond to humoral signals and distinct tissue-specific cues (2–4). Synchronization of clock circuits in the body ensures coordinated temporal orchestration of daily physiological processes. Disruption of clock synchrony, such as shiftwork or jet-lag, predispose to various disease risks including the development of cancer (5–7) and metabolic disorders (8–12). Better understanding of the external or endogenous stimuli that entrains the circadian clock provides fundamental knowledge of the mechanisms that underly environmental adaptation of the circadian clock (2).

The molecular circuit driving the ~24-hour rhythm of circadian oscillators is composed of a transcription-translation feedback loop (TTFL). Circadian Locomotor Output Kaput (CLOCK) and Brain and muscle Arnt-like 1 (Bmal1), the key transcriptional activators of this molecular clock feedback loop, heterodimerize and activate the transcription of clock repressor proteins, the Periods (Per1, Per2 & Per3) and Cryptochromes (Cry1 and Cry2). Ensuing cytosolic accumulation, phosphorylation and nuclear translocation of the Period and Cryptochrome proteins inhibit Bmal1/CLOCK-dependent transcription *via* direct interaction with the heterodimer. This negative transcriptional feedback cycle coupled with translational control constitutes the core clock regulatory loop. An additional Rev-erbα/ROR-mediated Bmal1 transcriptional oscillation re-enforces the robustness of this core clock mechanism (13).

Daily entrainment to environmental cues is essential for internal synchronization of the body clock system and adaptation to cyclic changes (2). Despite our current knowledge of the intricate molecular network driving clock oscillation and entrainment, how clock respond and adapt to its immediate niche environment is yet to be addressed. Various extracellular physical or chemical cues, including cell adhesion to extracellular matrix (ECM) *via* integrin or growth factor activation *via* cell-surface receptors, are transduced intracellularly by a signaling cascade that involves actin cytoskeleton remodeling and downstream transcriptional response mediated by serum response factor (SRF) in response to Myocardin-related Transcription Factor (MRTF) activation (14, 15). Integrins, through direct interactions with specific extracellular matrix (ECM) components, form focal adhesion complexes that connect the cellular physical environment with intracellular actin cytoskeleton network through a myriad of signaling pathways (16). Activation of integrin-mediated intracellular signaling transduction, including focal adhesion-associated kinase (FAK), Rho-GTPases, ROCK kinase and their associated effector molecules, transmits extracellular microenvironment cues to modulate actin polymerization *via* the formation of filamentous actin (F-actin) from monomeric globular actin (G-actin). Various growth factors or cytokines, including PDGF, TGF-β or Wnt, also elicit Rho-GTPase/ROCK signaling through cell surface receptors to engage actin cytoskeleton dynamic regulation. Actin polymerization leads to the release of Myocardin-related Transcription Factors (MRTF-A/B, MKL-1/2) from sequestration by G-actin monomers, with subsequent nuclear translocation and activation of SRF-mediated transcription (15). Nuclear shuttling of MRTF, in response to actin dynamic regulation, interacts with SRF on its cognate DNA binding motif, the CArG box, to control target gene expression (17). MRTF/SRF-controlled genes participate in various physiological processes during tissue development, growth and remodeling involving cytoskeleton organization, such as matrix adhesion, migration, proliferation and differentiation (14, 18–20).

Serum is known as an universal synchronizing signal of cellular clocks (21). Notably, serum-stimulated intracellular actin turnover entrains the liver clock (22, 23). However, limited studies so far directly address the effect of actin cytoskeleton remodeling and its interaction with ECM on clock properties, particularly pertaining to the signaling cascades involved in modulating MRTF/SRF activity. In the current study, we employed pharmacological and genetic approaches targeting specific steps of integrin-actin cytoskeleton-MRTF/SRF pathway to establish its role in modulating clock oscillation. Our study revealed that this cascade linking cellular niche cues with intracellular actin dynamics impacts key functions of the circadian clock.

## Results

### Pharmacological perturbations of actin polymerization modulate clock oscillation

Polymerization of monomeric actin, in response to upstream signals, is the key regulatory step that releases MRTF from G-actin sequester to activate SRF-mediated transcription (14). To test whether actin dynamic-associated signaling modulates circadian clock, we first determined the effects of pharmacological agents that disrupt or promote actin polymerization, using mouse fibroblasts containing Per-2 promoter-driven luciferase reporter knock-in (Per2-Luc) (24). Monitoring of Per2-Luc fibroblast bioluminescence activity revealed that Cytochalasin D (Cyt D), a known actin depolymerization compound (25, 26), significantly altered clock cycling properties, as shown by the average tracing at varying concentrations (Fig. 1A & 1B). Cyt D induced a dose-dependent increase of oscillation amplitude (Fig. 1C), with concomitant lengthening of period (Fig. 1D). Cyt D treated-cells also displayed consistent phase advance as compared to controls within the tested range of 1uM to 5uM (Fig. 1E). To determine the effect of Cyt D on intracellular actin organization, we performed immunofluorescence staining of F-actin stress fibers by phalloidin. Cty D-treated cells display marked reductions of F-actin that persisted at 24 hours after treatment, with cells viable at both concentrations tested (Fig. 1F). The loss of actin polymerization is also consistent with the gradual change in rounded up cell morphology observed (Suppl. Fig. S1). As expected of Cyt D action in interfering with actin polymerization and inhibiting MRTF/SRF activity, expression of known MRTF/SRF target genes, connective tissue growth factor (*Ctgf*) and Four and half LIM-domain protein 1 (*Fhl1*), were markedly attenuated by Cyt D (Fig. 1G). *Mkl1* transcript was also reduced, although *Srf* expression was not altered. Analysis of core clock genes revealed significant effects of Cyt D on inhibiting *CLOCK, Per2* and *Cry2* expression, without altering other clock circuit components (Fig. 1H). Suppressed clock gene expression by Cyt D suggests potential transcriptional regulation induced by actin dynamic-related signaling, and dampening of positive clock component with augmented negative regulatory loop may contribute to the period lengthening effect of Cyt D.

**Figure 1.**
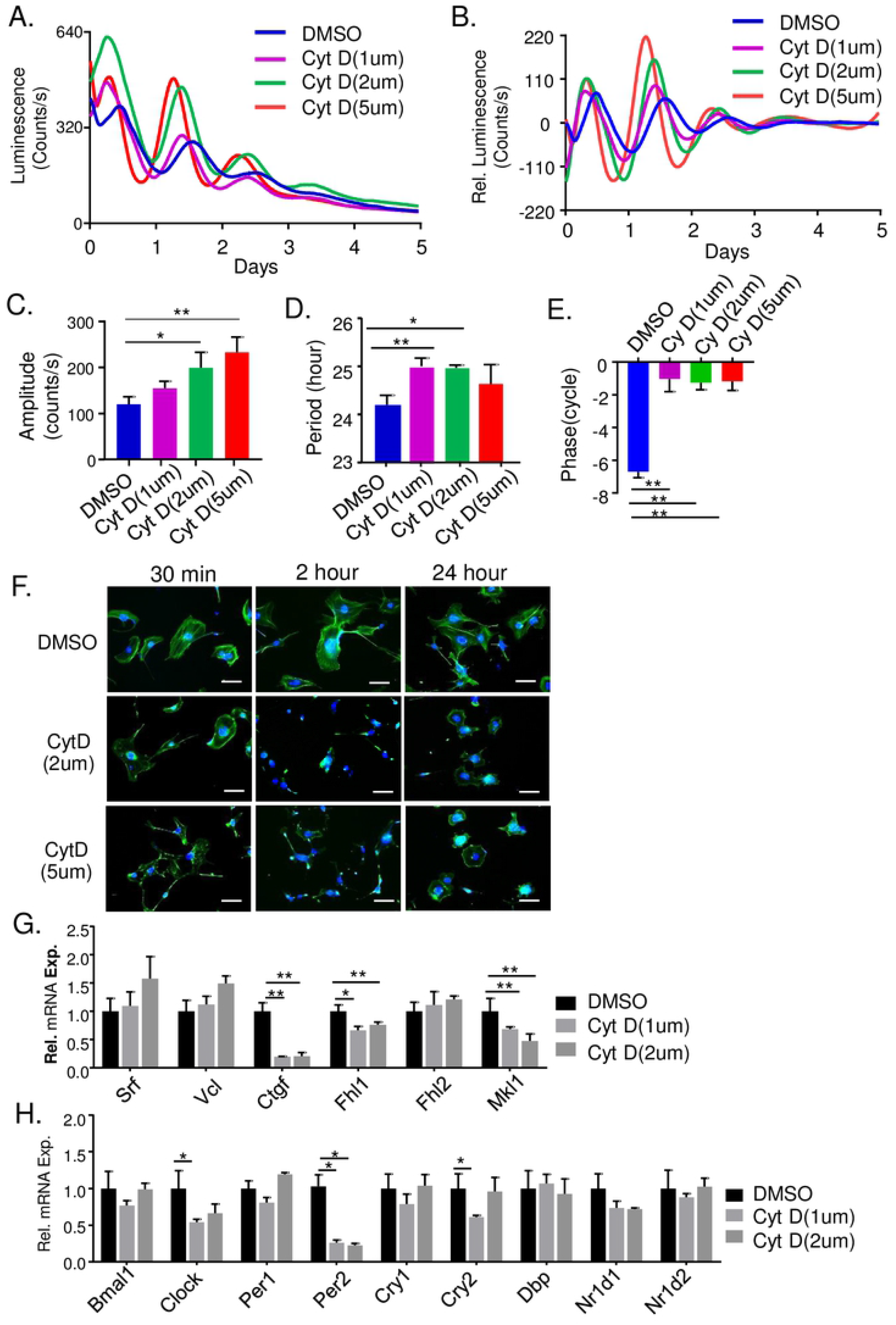
Effect of actin-depolymerizing chemical Cytochalasin D on modulating clock rhythm. (A, B). The original (A), and baseline-subtracted (B) bioluminescence plots of Per2-Luciferase knock-in fibroblasts subjected to Cytochalasin D (Cyt D) treatment at indicated concentration. (C-E) Quantitative analysis of clock cycling properties, including amplitude (C), period length (D), and phase angle (E, n=3). (F) Representative fluorescence images of F-actin stress fibers by phalloidin staining in control (DMSO) and Cyt D-treated Per-2 knock-in fibroblasts. Scale bar: 100 μm. (G & H) RT-qPCR analysis of Cyt D effect on expression of target genes and signaling components of MRTF/SRF-mediated transcription (G), and core clock genes (H, n=3). *, **: P≤0.05 or 0.01 Cyt D vs. DMSO.

We next examined whether another molecule that inhibits actin polymerization, latrunculin B (Lat B), display shared clock-modulatory activities with Cyt D (27). Indeed, Lat B led to a similar dose-dependent effect on increasing clock amplitude (Fig. 2A–2C), together with significant period lengthening at 2 uM (Fig. 2D). In contrast, Lat B induced a stronger effect on phase advance as compared to that of Cyt D (Fig. 2E). The loss of F-actin organization induced by Lat B was also more rapid and robust than that of Cyt D, with substantially reduced phalloidin staining apparent at 30 minutes following treatment and persisted throughout the time course examined (Fig. 2F). Lat B on actin cytoskeleton is further reflected in the rapid change of cell morphology (Suppl. Fig. S1). Notably, Lat B displayed marked inhibition of SRF target genes at concentrations tested, with near complete inhibition of *Ctgf* and strong suppression of *Srf, vinculin* (*Vcl*), *Fhl1* and *Fhl2* (Fig. 2G). Interestingly, Lat B induced ~70% suppression of *Bmal1* expression, while *Per1, Dbp* and *Nr1d2* were induced Lat B at higher concentration (Fig. 2H).

**Figure 2.**
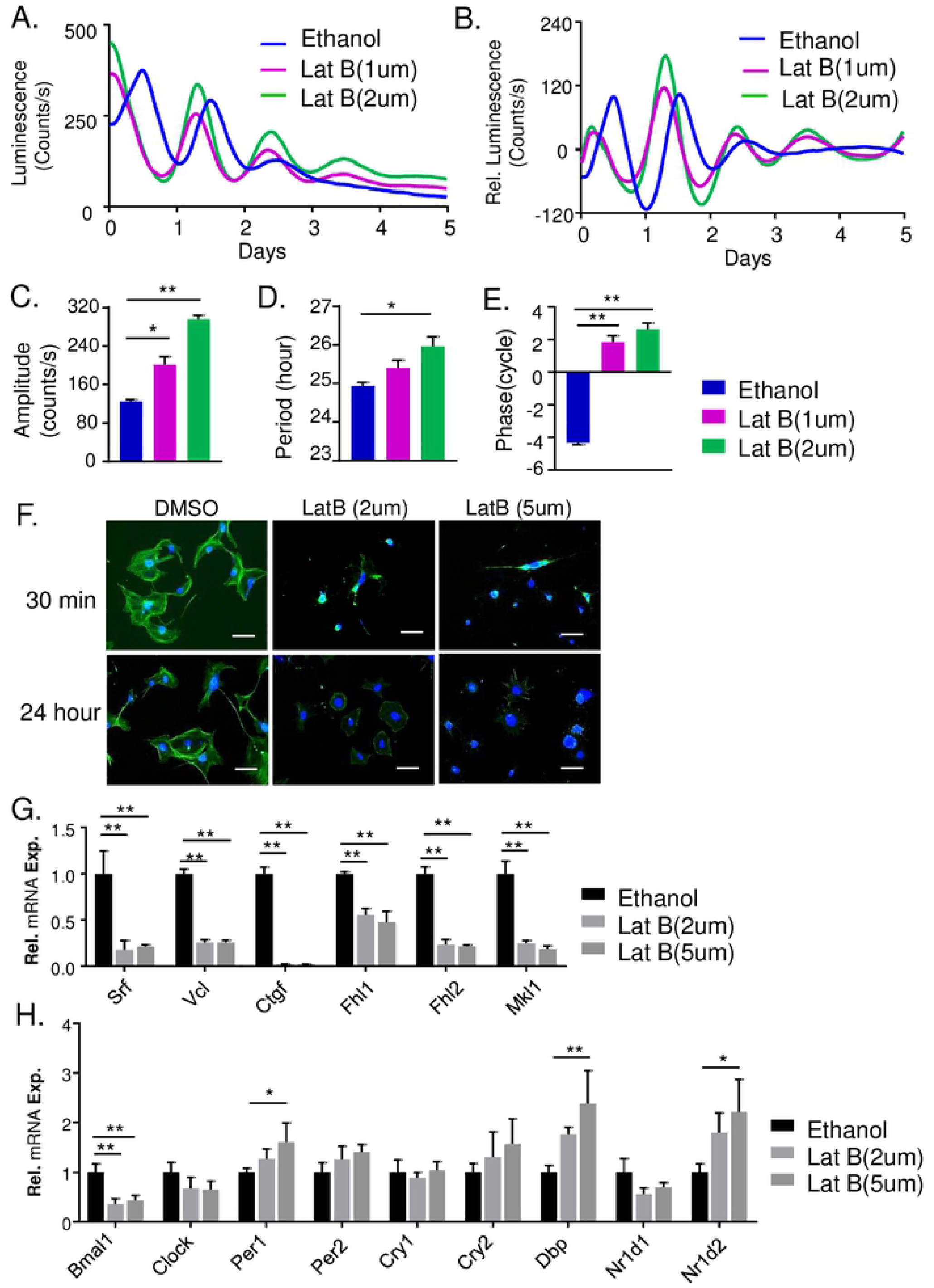
Effect of Latrunculin B on clock rhythm modulation. (A, B). The original (A), and baseline-subtracted plots (B) of Per2-Luciferase knock-in fibroblasts bioluminescence subjected to Latrunculin B (Lat B) treatment at indicated concentrations. (C-E) Quantitative analysis of clock cycling properties, the amplitude (C), period length (D, n=3), and phase angle in response to Lat B (E). (F) Representative fluorescence images of F-actin by phalloidin staining in control (DMSO) and Lat B-treated Per-2 knock-in fibroblasts. Scale bar: 100 μm. (G, H) RT-qPCR analysis of the effect of Lat B expression of MRTF/SRF-related signaling pathways (G), and core clock genes (H, n=3). *, **: P≤0.05 or 0.01 Lat B vs. Ethanol.

Furthermore, we tested whether promoting actin polymerization *via* Jasplakinolide (Jas) (27), thereby stimulating MRTF/SRF-mediated transcription, affects clock function. In contrast to Cyt D, subjecting Per2-Luc fibroblasts to Jas significantly decreased clock amplitude at 0.5 uM (Fig. 3A–C). In addition, Jas treatment resulted in dose-dependent period shortening from 0.1 to 0.5 uM concentration (Fig. 3D), with only significant effect on phase delay at 0.5 uM (Fig. 3E). Jas induced only moderate increase in F-actin immunofluorescence as shown in Fig. 3F, likely due to the well-organized robust F-actin structure present in fibroblasts under normal conditions. Gene expression analysis confirmed Jas induction of SRF-mediated transcription activation, as indicted by marked up-regulations of SRF target genes including *Vcl, Ctgf, Fhl1, Fhl2* together with *Srf* itself (Fig. 3G). In comparison to the clock gene repression induced by Cyt D, Jas elicited wide-spread inductions of clock genes examined, including *Bmal1, Per2, Cry2, Nr1d1* and *Nr1d2* in a dose-dependent manner (Fig. 3H), although *Per1* level was reduced.

**Figure 3.**
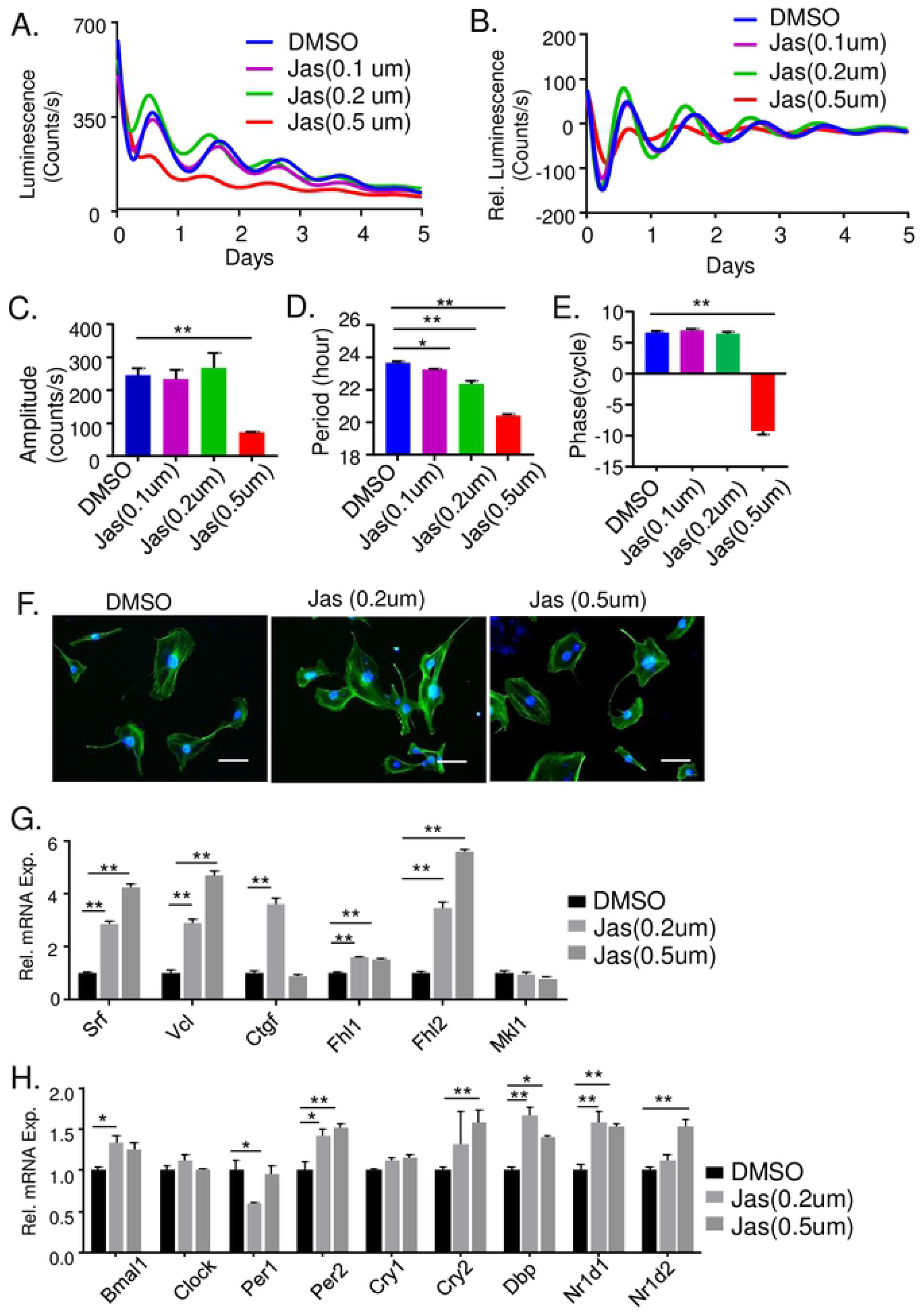
Effect of actin-polymerizing Jasplakinolide on clock modulation. (A, B) Raw bioluminescence (A), and baseline-subtracted plots (B), of Jasplakinolide (Jas) effect on Per2-Luciferase bioluminescence activity in mouse fibroblasts at 0.1 to 0.5 uM concentration. (C-E) Quantitative analysis of clock cycling properties, the amplitude (C), period length (D) and phase angle (E, n=3) in response to Jas. (F) Representative fluorescence images of F-actin by phalloidin staining in control (DMSO) and Jas-treated Per-2 knock-in fibroblasts for 2 hours. Scale bar: 100 μm. (G & H) RT-qPCR analysis of the effect of Jas on expression of MRTF/SqRF signaling components (G), and core clock genes (H, n=3). *, **: P≤0.05 or 0.01 Jas vs. DMSO.

### Genetic inhibition of SRF or MRTF function modulates clock property

Intracellular signaling induced by actin dynamics ultimately results in MRTF/SRF-mediated transcription activation response. Using siRNA-mediated silencing of these effectors of actin dynamics, we tested whether SRF or MRTF-A mediates the effects of actin cytoskeleton-modifying molecules on clock modulation. One of the three *Srf* siRNA tested was most effective in abolishing SRF protein expression (Fig. 4A). As shown in Fig. 4B, *Srf* silencing altered clock oscillatory properties similarly as actin-depolymerizing agents, leading to increased period length with a tendency toward higher amplitude (Fig. 4C & 4D). We nest performed serum shock-induced synchronization to determine the effect of SRF inhibition on clock gene expression at two circadian times, CT12 (CT: circadian time) and CT24. *Mkl1, Fhl1* and *Fhl2* displayed circadian time-dependent regulations with higher expressions at CT24 than that of CT12 (Fig. 4E). Knockdown of *Srf* markedly reduced *Ctgf* and *Fhl1* expression at both time points, as compared to that of the scrambled controls (SC), while *Mkl1* and *Fhl2* levels were significantly lower only at CT24. Analysis of clock genes revealed inductions of *Per1* and *Cry1* at CT24 than that of CT12, while both were significantly down-regulated by *siSrf* at CT24 (Fig. 4F). Furthermore, loss of *MRTF-A* by siRNA knockdown (*siMrtf*) resulted in similar effects as SRF inhibition. Two *siMrtf* examined largely abolished MRTF-A protein expression (Fig. 4G). Similar to *Srf* inhibition, *Mrtf* silencing resulted in augmented clock oscillation amplitude (Fig. 4H & 4I) with increased period length (Fig. 4J), although degree of the effects differing between two knockdowns. One *siMrtf* used induced a significant phase advance (Fig. 4K). Together, these findings suggest that SRF and MRTF are directly involved in clock oscillation and exert transcriptional regulation on core clock circuit.

**Figure 4.**
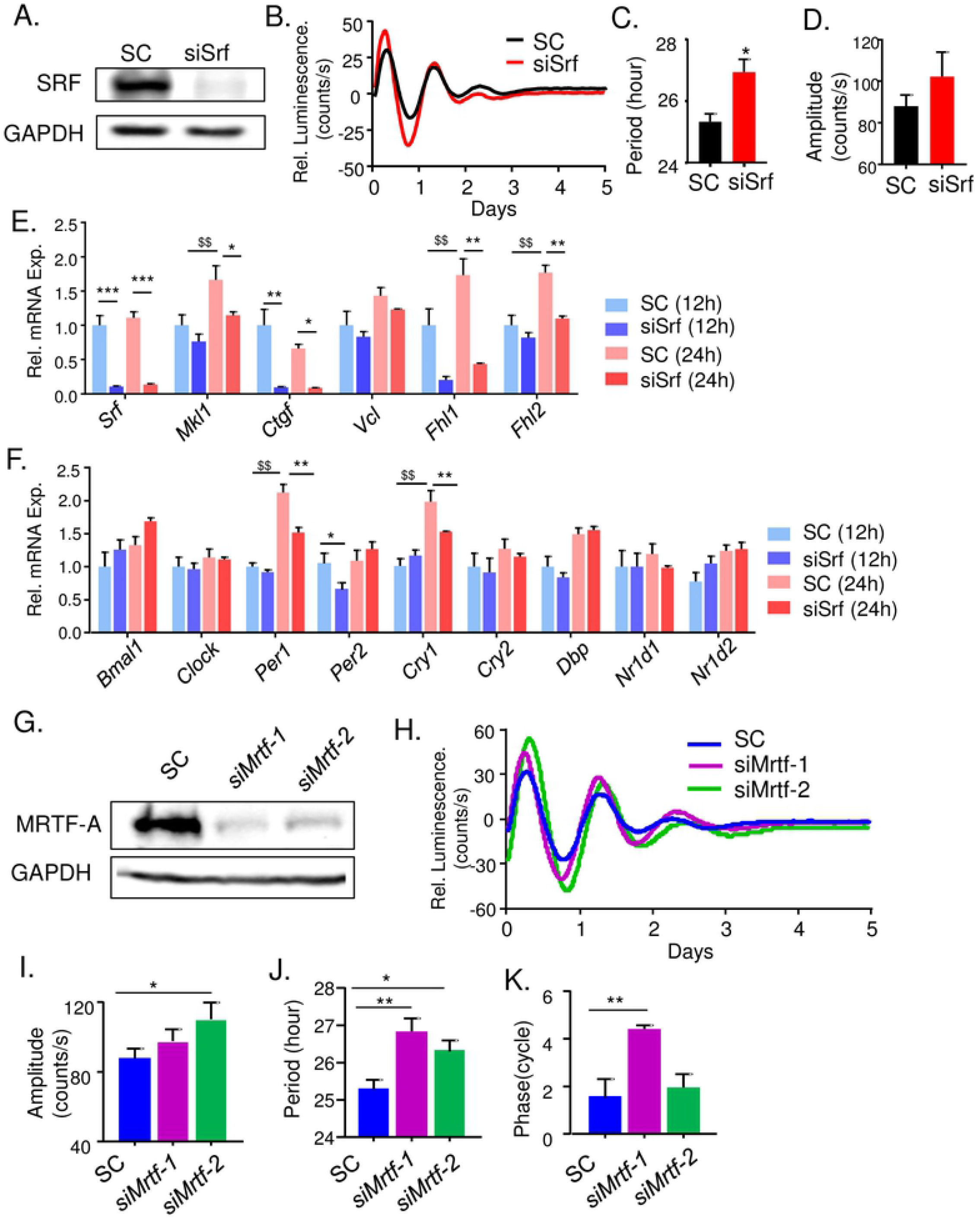
Genetic loss-of-functions of SRF and MRTF-A influence clock function. (A-F). Effect of *Srf* silencing on core clock properties. (A) Immunoblot analysis of SRF protein expression in Per2-Luc fibroblasts with transient knockdown of scrambled control (siSC) or *siSrf*. (B-D) Representative bioluminescence tracings of Per2-Luc fibroblasts (B) with quantification of clock amplitude (C) and period length (D, n=3). *, **: P≤0.05 or 0.01 *siSrf* vs. SC. (E & F) RT-qPCR analysis of MRTF/SRF-related signaling components (E), and core clock gene expression (F) at 12 and 24 hours after serum shock synchronization (n=3). $, $$: P≤ 0.05 or 0.01 control 24hr vs. 12hr; and *, **: P≤0.05 or 0.01 *siSrf* vs. SC. (G-K) Effect of MRTF-A silencing on clock activity. Immunoblot analysis of MRTF-A protein level (G), representative bioluminescence of Per2-Luc fibroblasts transfected with scrambled control (SC) or *siMrtf* (H), and quantification of amplitude (I), clock period (I), and phase angle (K, n=3). *, **: P≤0.05 or 0.01 *siMrtf* vs. SC.

### Inhibition of upstream signaling pathways inducing actin cytoskeleton remodeling modulates clock function

The Rho GTPases (RhoA, Rac1 and Cdc42) and Rho-associated kinase (ROCK) are upstream signaling molecules transmitting cell surface stimuli to modulate actin dynamics (25, 26). Cell surface ECM or biochemical cues, *via* integrin or receptor tyrosine kinases, activate Rho GEFs and Rho/ROCK-mediated kinase cascade to alter G-actin to F-actin ratio (23). Using a specific ROCK kinase inhibitor, Y27632, we determined the effect of Rho GTPase signaling perturbation upstream of actin dynamics on circadian clock. As shown in Fig. 5A–5E, Y27632 exerted similar effects on clock properties as actin-disrupting molecules, with significantly increased cycling amplitude (Fig. 5A–C), prolonged period length (Fig. 5D) and phase advance (Fig. 5E). Y27632 induced a robust, somewhat delayed effect on F-actin as compared to actin depolymerizing agents, as indicated by the cell shape change and decreased phalloidin staining most evident at 2 hours after administration (Suppl. Fig. S2). As expected with inhibition of MRTF/SRF activity, Y27632 markedly down-regulated SRF target genes with nearly ~80% lower mRNA transcripts of *Srf, Vcl, Ctgf* and *Mkl1*, as compared to DMSO-treated controls (Fig. 5F). Y27632 also suppressed *Per2, Nr1d1* and *Nr1d2* expression similar to that of Cyt D effect (Fig. 5G), whereas its induction of *Cry1* and *Cry2* are distinct. In response to ROCK activation, myosin light chain kinase (MLCK) phosphorylation promotes F-actin formation and actomyosin contractility involved in cellular migration (28). Using ML7, a specific MLCK inhibitor, we tested whether this MLCK activation downstream to ROCK modulates clock function (29, 30). ML7 induced significantly increased clock amplitude (Fig. 5H & 5I) with phase advance at 20 uM (Fig. 5K), similar to actin-depolymerizing molecules, although it reduced period length (Fig. 5J). Gene expression analysis revealed that ML7 consistently attenuated SRF target gene expression (Fig. 5L), as expected, whereas its effects on clock genes were limited to inhibition of *Per2* with up-regulation of *Cry2* (Fig. 5M). Thus, perturbing signaling events up-stream of actin cytoskeleton remodeling, to a large extent, recapitulated the effects of actin depolymerization on clock modulation.

**Figure 5.**
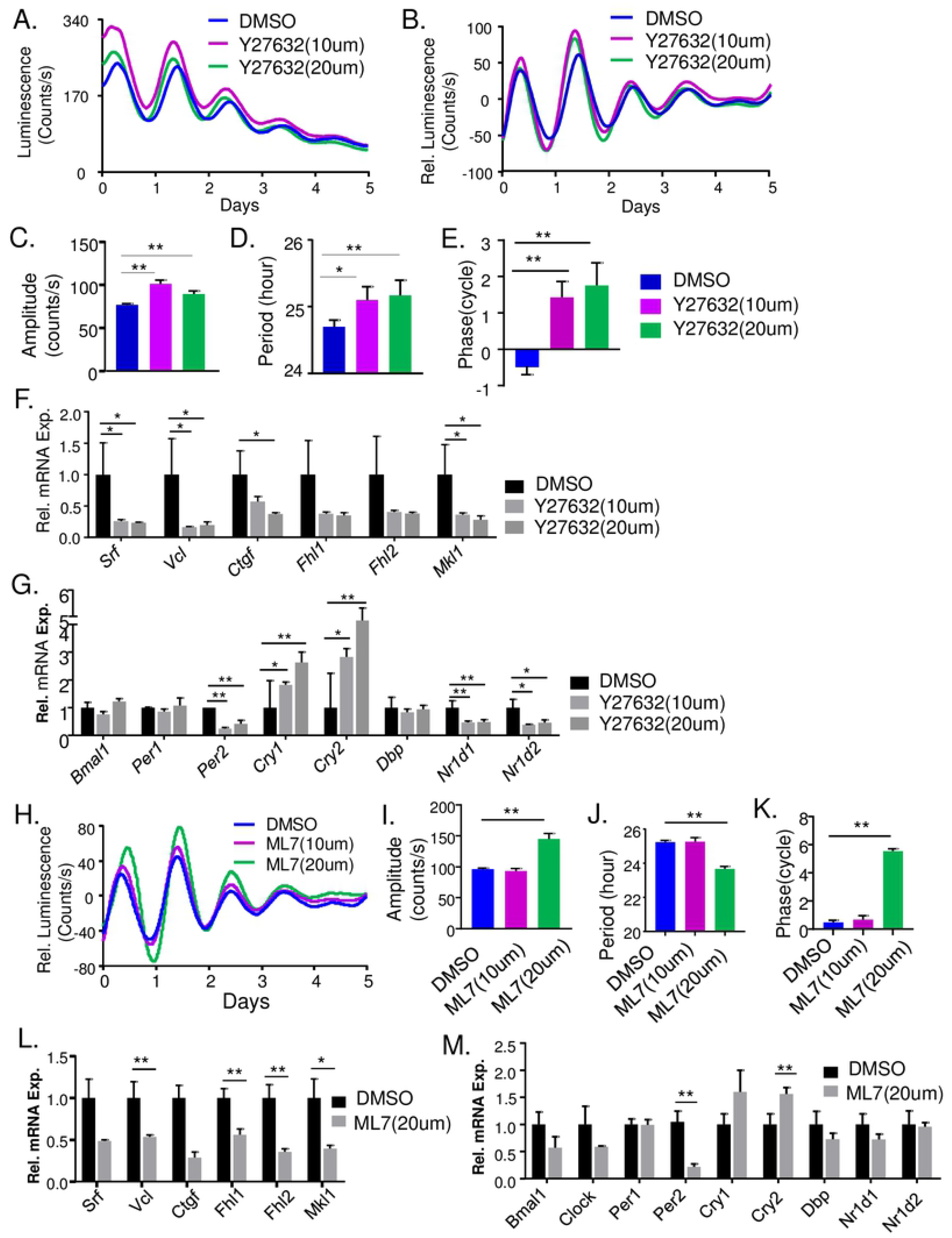
Effect of ROCK inhibition by Y27632 and MLCK inhibition by ML7 on circadian clock properties. (A-E). Representative raw bioluminescence (A), and baseline-subtracted plots (B) of dose-dependent effects of ROCK inhibitor Y27632 on Per2-Luciferase activity, with quantitative analysis of clock cycling amplitude (C), period length (D), and phase (E, n=3). (F & G) RT-qPCR analysis of the effect of Y27632 on MRTF/SRF signaling pathway (F), and core clock genes (G, n=3). (H-M) Effect of myosin-light chain kinase inhibitor ML7 on clock rhythm. (H-K) ML7 effect on baseline-subtracted bioluminescence plot of Per2-Luciferase (H), with quantitative analysis of amplitude (I), period length (J) and phase (K, n=3). (L & M) RT-qPCR analysis of ML7 regulation of MRTF/SRF signaling pathways (L), and core clock genes (M). *, **: P≤0.05 or 0.01 Y27632 or ML7 vs. DMSO control.

### Integrin-mediated extracellular matrix adhesion signaling modulates clock function

ECM constitutes the immediate physical micro-environment cells residing in, and integrin-mediated focal adhesion complex links ECM components with intracellular actin cytoskeleton (14). Focal adhesion kinase (FAK), together with additional components of the focal adhesion complex, transduces ECM cues to modulate F-actin stress fiber formation with activation of MRTF-SRF transcriptional response (16). Based on the finding that actin dynamic-induced MRTF/SRF activity modulates circadian clock, we postulated that blocking integrin-associated focal adhesion signaling may impact clock function. We thus determined the effect of integrin αV-targeting cyclic peptide (cRGD) to block integrin interaction with ECM, as compared to a scrambled sequence peptide control. As shown in Fig. 6A & 6B, cRGD-treated cells displayed significantly higher oscillation amplitude at 2uM, with a tendency toward longer period (Fig. 6C) and significant phase advance (Fig. 6D). The effect of integrin blocking peptide was confined to a narrow molar range, as 5uM completely abolished clock cycling, suggesting tight stoichiometry of RGD action in blocking integrin binding sites involved in cell-matrix interactions. RT-qPCR analysis demonstrated down-regulations of SRF transcriptional targets, *Srf, Vcl* and *Ctgf*, indicative of inhibition of MRTF/SRF signaling by cyclo-RGD (Fig. 6E), together with significantly suppressed *Bmal1* and *Per2* as compared to control peptide-treated cells (Fig. 6F). Furthermore, we tested whether inhibiting FAK that mediates integrin-induced signaling affects clock properties using genetic knockdown. Three *siFAK* were found to markedly reduce FAK protein level (Fig. 6G), and the silencing of FAK had significant effects on modulating clock activity (Fig. 6H). Although *siFAK*s did not alter cycling amplitude (Fig. 6I), these siRNAs induced consistently prolonged clock period (Fig. 6J) with significant phase advance (Fig. 6K).

**Figure 6.**
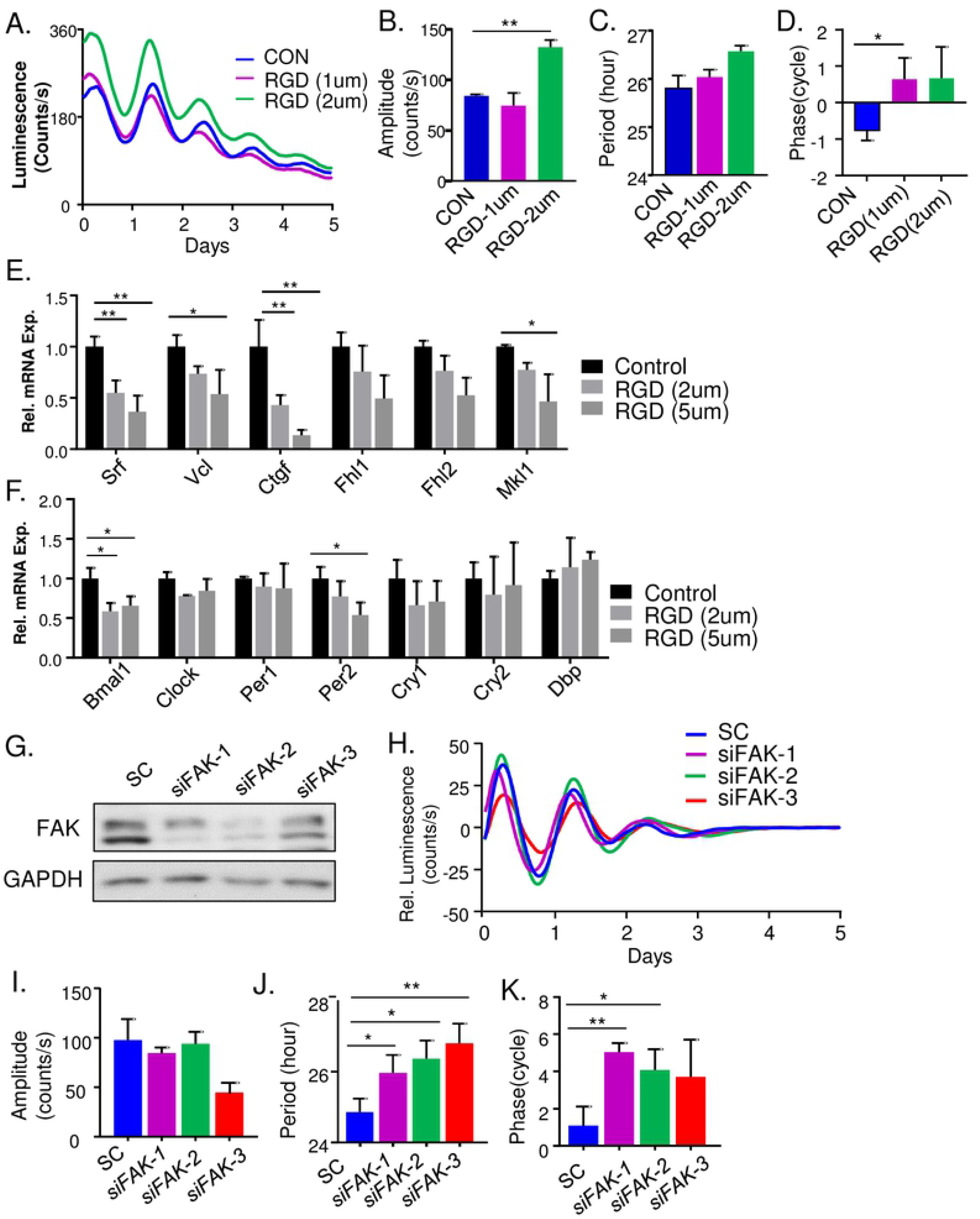
Inhibition of integrin-mediated focal adhesion signaling modulates clock function. (A-E) Effect of integrin blocking peptide Cyclo RGD on clock function. Representative bioluminescence of dose-dependent effects of RGD on Per2-Luciferase activity (A), with quantitative analysis of clock cycling amplitude (B), period length (C) and phase (D, n=3). (E & F) RT-qPCR analysis of the effect of integrin blocking peptide on MRTF/SRF signaling components (E), and core clock gene expression (F, n=3). *, **: P≤0.05 or 0.01 RGD blocking peptide vs. control peptide. (J-K) Effect of focal adhesion kinase (FAK) gene silencing by siRNA knockdown on clock oscillation. Immunoblot analysis of FAK protein level (G), representative bioluminescence of Per2-Luc fibroblasts transfected with scrambled control (SC) or *siFAK* (H), with quantification of clock amplitude (I), period length (J) and phase (K, n=3). *, **: P≤0.05 or 0.01 *siFAK* vs. SC.

### MRTF and SRF exerts direct transcriptional control of core clock regulators

In response to altered G-actin/F-actin ratio, MRTF translocates to the nucleus and binds to SRF to activate gene transcription *via* the CArG box motif (31). While SRF association with cognate DNA binding sites is largely constitutive, MRTF is responsive to extracellular stimuli-elicited Rho-ROCK-actin signaling (18, 23). To test actin dynamic-induced MRTF-SRF activity exerts direct transcriptional control of the molecular clock circuit, we determined MRTF or SRF chromatin occupancy on core clock genes. Consensus MRTF/SRF DNA binding element, CArG box, were identified within gene regulatory regions containing proximal promoter (+-2kb of transcription start site) of core clock genes using TRANSFAC. Using chromatin immunoprecipitation-qPCR (ChIP-qPCR), ~8-10-fold of SRF enrichment on known CArG sites within *alpha-actin* and *vinculin* promoters were detected over IgG control, as expected (Fig. 7A). Similar degrees of ~6-10-fold enrichment of SRF occupancy were found on clock gene regulatory regions within *Per1, Per2, Nr1d1*, and *Nfil3*, demonstrating these clock components as direct SRF target genes. We next performed immunoprecipitation with a MRTF-A antibody to further determine MRTF/SRF-mediated transcriptional control of these genes. As shown in Fig. 7B, we observed similar extent of MRTF-A chromatin association of the clock gene regulatory regions examined. Interestingly, MRTF-A occupancy of identified CArG site on Per2 promoter was markedly higher than that of SRF. To explore whether loss of SRF or MRTF affects their chromatin occupancy on clock target genes, we generated stable cell lines containing SRF or MRTF-A/B shRNA knockdown (32). Loss of *Srf* by stable knockdown in Per2-Luc fibroblasts largely abolished binding to identified regulatory regions of clock gene targets, with detected enrichment comparable to IgG control (Fig. 7C). Similar loss of MRTF-A binding to target promoters were observed in cells with stable expression of MRTF-A/B shRNA (Fig. 7D).

**Figure 7.**
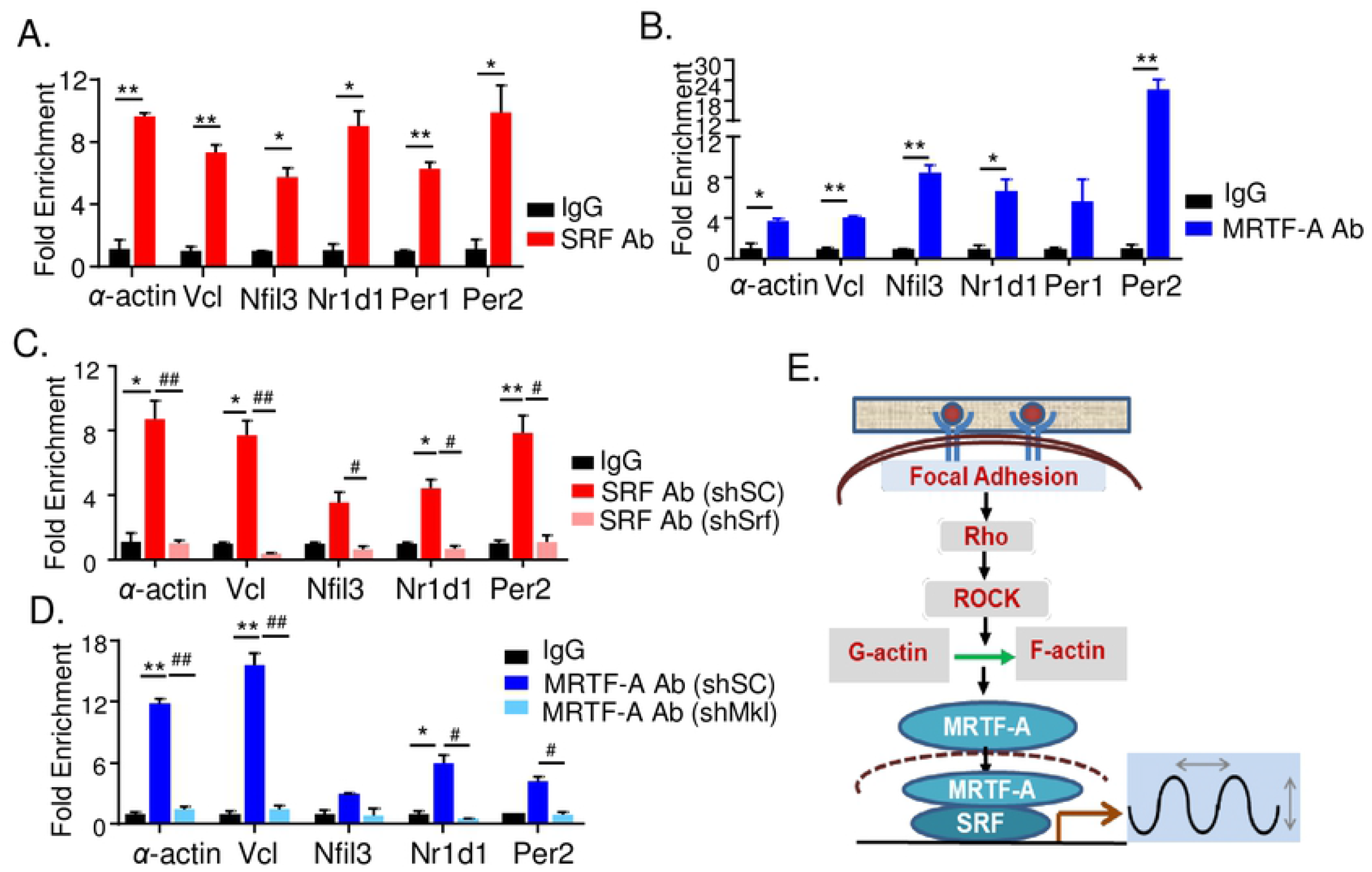
Direct MRTF/SRF transcriptional control of core clock genes. (A, B) ChIP-qPCR analysis of chromatin occupancy of SRF (A), and MRTF-A (B), on regulatory regions of core clock genes, as compared to IgG control. SRF and MRTF-A binding to known CArG sites of α-actin and vinculin (Vcl) promoters were as positive controls. (C, D) ChIP-qPCR analysis of SRF (C), and MRTF (D) chromatin occupancy in fibroblasts with stable expression of scrambled control (*shSC*), *Srf* shRNA knockdown (*shSrf*, C), or *Mrtf* shRNA knockdown (*shMrtf*, D). *, **: P≤0.05 or 0.01 SRF or MRTF-A antibody vs. IgG control. #, ##: P≤0.05 or 0.01 *shSrf* or *shMrtf* vs. *shSC*. (E) Schematic model of focal adhesion-actin cytoskeleton-MRTF/SRF signaling cascade in transducing cellular niche cues to circadian clock circuit.

Notably, the extent of MRTF-A enrichment was mostly attenuated in cells with scramble control expression as compared to normal Per2-Luc fibroblasts. Together, these findings identified direct MRTF/SRF transcriptional control within specific core clock components. Collectively, our results revealed a MRTF/SRF-mediated regulatory mechanism in controlling clock gene transcription, and this mechanism is responsive to extracellular niche signals transmitted *via* the Rho-ROCK-actin remodeling signaling cascade (Fig. 7E).

## Discussion

Employing pharmacological perturbation with genetic approaches, we investigated the effect of integrin-actin cytoskeleton-MRTF/SRF signaling on circadian clock modulation. Our study established the role of distinct steps of this signaling transduction cascade in determining clock activity, implicating this pathway in transducing extracellular physical niche signals to entrain circadian clock. Given the importance of actin cytoskeleton-MRTF/SRF signaling in driving cellular adhesion, migration, proliferation and differentiation, circadian clock outputs could be intimately linked with these fundamental aspects of cellular behavior involved in tissue development, growth and remodeling (14, 18, 20).

Our results obtained from interfering with actin cytoskeleton organization and consequently, the modulation of MRTF/SRF activity, yielded largely consistent effects on clock oscillatory amplitude and period length. Chemicals disrupting actin polymerization, including cytochalasin D and latrunculin B, augmented clock cycling amplitude with longer period, whereas facilitating F-actin polymerization through Jasplakinolide led to opposite effects of reduced amplitude with shortened period. Blocking signaling up-stream of actin cytoskeleton remodeling by ROCK or MLCK inhibitors also demonstrated increased oscillation amplitude and period length. Higher clock amplitude as a result of genetic inhibition of *Srf* or *Mrtf-a*, the transcription transducers of actin dynamics-induced response, further corroborated findings from pharmacological perturbations of actin dynamics by demonstrating the direct effects of *Srf* and *Mrtf-a*. These perturbations robustly influence clock gene transcription consistent with MRTF/SRF signaling activity, and analysis of putative clock gene promoters identified Per1, Per2, Cry2 and Nfil3 as direct transcriptional targets of SRF and MRTF.

We observed certain divergent effects among distinct pharmacological interventions of the integrin-actin cytoskeleton-MRTF/SRF cascade on clock activity. ML7 inhibition of MLCK led to period shortening, potentially due to off-target effects on inhibiting kinases other than MLCK (29, 30). Despite consistent modulatory effects on negative clock regulators, including Per1, Per2 and Nr1d1, specific gene expression profiles of clock components induced by actin polymerizing chemicals differs. The precise regulation of clock properties by these molecules may depend on the distinct combined effects of clock transcriptional regulation involving the molecular transcriptional-translational feedback loop. Interestingly, the effect of blocking integrin-mediated adhesion signaling on clock, by RGD or FAK knockdown, are relatively moderate as compared to actin-disrupting molecules. RGD had significant effect on amplitude, while FAK silencing increased period length. A multitude of intracellular molecules and pathways mediates transduction of extracellular matrix cues to actin dynamic-driven MRTF/SRF transactivation (14). Thus, it is conceivable that interfering with cell-matrix interactions through RGD or FAK only may not be as robust as modulating the shared downstream signaling through actin cytoskeleton-MRTF/SRF activity.

As a time-keeping mechanism to anticipate and adapt to cyclic environmental stimuli, circadian clock can be entrained by various extrinsic and endogenous timing cues (2). Serum stimulation is recognized as a strong entrainment signal for peripheral clocks (21). Gerber et al. discovered that serum-induced actin remodeling and modulation of MRTF/SRF activity in liver mediates clock entrainment (22), and this pathway is responsive to upstream signaling events involving Rho kinase activation in fibroblasts (23). Our study elucidates how actin cytoskeleton-associated signaling nodes transduce integrin-associated niche cues to MRTF/SRF-mediated transcriptional events that control clock gene expression. These findings implicate a sensing mechanism of circadian clock of its immediate extracellular microenvironment, which may facilitate cellular adaptation to its physical niche. Circadian clock is known to influence stem cell behaviors in distinct tissue remodeling processes involving epidermal stem cells, intestinal stem cells or myogenic progenitors (33–38). Our finding of the mechanistic connection of circadian clock modulation by its physical niche implicates its potential contribution to certain stem cell processes during tissue development or remodeling that involves cellular interaction with its niche environment (39).

Within a myriad of stem cell compartments, proper circadian clock functions are required for tissue growth or remodeling by coordinating stem cell behavior with environmental stimuli (39). Clock oscillation is known to modulate epidermal stem cell activation status that determines its responsiveness to epidermal remodeling cycles (33). Hair follicle stem cell activation was gated by circadian clock to protect against genotoxic stress during regeneration (34), while intestinal stem cell regeneration was found to be regulated by clock genes in Drosophila or murine models (35, 40). Our previous studies demonstrated that clock regulators, Bmal1 and Rev-erbα, plays antagonistic roles in muscle regeneration by modulating muscle stem cell proliferation and differentiation in response to injury (36–38, 41). Stem cell quiescence and self-renewal properties are intimately linked with its physical niche environment (42). Thus, it will be intriguing to explore how clock modulation by ECM microenvironment may apply to stem cell biology. Furthermore, cell-cell interactions such as those mediated by adherens junctions also require actin cytoskeleton anchoring, while intercellular couplings are known to promote the robustness of clock oscillation (43). It is thus possible that circadian oscillation may respond to intercellular interactions that are transduced to actin cytoskeleton organization. The intriguing interplay between cellular physical niche signals and timing system may direct proper developmental progressions with appropriate spatial and temporal coordination.

Integrins bridges extracellular matrix with intracellular actin cytoskeleton (16, 44). Integrin-linked actin remodeling thus transduces cellular physical niche environment through the MRTF/SRF signaling cascade (15). Our findings linking integrin-associate signaling with clock modulation suggests that integrin may transduce specific physical niche cues from extracellular matrix to induce clock synchronization. It is thus plausible that physical properties of ECM, including stiffness or composition, may influence clock functions in distinct cell types to modulate its biological output pathways. Interestingly, Yang et al. demonstrated that matrix stiffness, but not composition, strongly modulates clock oscillation in mammary epithelial cells (45), and a related study revealed that softer 3-D matrix induced stronger clock gene oscillations than stiff 2-D matrices (46). Our findings are in line with these observations from mammary epithelial cell clock, although these studies did not address the role of MRTF/SRF-mediated transcriptional control in clock regulation. While we found that cytoskeleton disruption augmented clock oscillation in Per2-Luc fibroblasts, agents promoting actin de-polymerization or ROCK kinase inhibition by Y27632 similarly induced clock amplitude in mammary epithelial cells (45). In contrast, fibroblasts isolated from mesenchyme of lung and mammary tissue were reported to display inverse regulation by matrix rigidity (46), suggesting complex tissue matrix-specific regulations of clocks in distinct cell types (47). Potential cell type-specific interactions of cells with its extracellular niche and precise regulations of the Rho-ROCK-actin signaling cascade warrants further examination in diverse cellular or *in vivo* models.

As our study revealed, small molecule modulators of actin dynamic and related signaling molecules significantly impact circadian clock function, mediated by MRTF/SRF transcriptional activity. These agents, such as RhoA/ROCK inhibitors or SRF modulators, may have unintended clock effects that are implicated in their biological activity. On the other hand, these pharmacological modulators or their chemical derivatives may have potential disease applications. With the recent interests in targeting clock for cancer and metabolic diseases (48–53), our finding of integrin-actin cytoskeleton-MRTF/SRF-mediated modulation of circadian clock properties could be applicable for disease therapies that warrants future studies.

## Materials and methods

### Cell Culture

Mouse primary fibroblasts originally obtained from enzymatic digestion of tail of mPER2::LUC-SV40 knock-in mice the mPer2 Luciferase-SV40 reporter (54) that overcame replicative senescence. These cells and U2OS cell line containing Bmal1-Luciferase or Per2-Luciferase reporter were kind gifts from Dr. Steve Kay at University of Southern California. Cells were maintained, in DMEM supplemented with 10% fetal bovine serum and antibiotics in a standard tissue culture incubator at 37°C with 5% CO2, as described previously (55). Cells were seeded in 24-well culture plates, grown to confluence, and switched to Lumicycle bioluminescence recording media for continuous Lumicycle recording for 7 days. Lumicycle recording media contains HEPES-buffered, air-equilibrated DMEM with 10 mM HEPES, 1.2 g/L NaHCO3, 25 U/ml penicillin, 25 g/ml streptomycin, 2% B-27, and 1 mM luciferin.

### Chemicals and Reagents

Cytochalasin D, Latrunculin B, Jasplakinolide, ML7 and Rock inhibitor Y27632 were purchased from Cayman Chemicals. Integrin RGD blocking peptide Cyclo [Arg-Gly-Asp-D-Phe-Val] and control Cyclo [Arg-Ala-Asp-D-Phe-Val] were obtained from Enzo Life Sciences.

### Luminometry, Bioluminescence Recording and Data Analysis

Measurement of bioluminescence rhythms from Per2-Luc mouse fibroblast cell culture were conducted using a luminometer LumiCycle 96 (Actimetrics), as described (53, 55, 56). Briefly, 24-well tissue culture plates were sealed with plastic film and placed inside a bacterial incubator maintained at 36°C, 0% CO2. Luminescence from each well was measured for ~70 s at intervals of 10 min and recorded as counts/second. 7 days of real-time bioluminescence recording data was analyzed using LumiCycle Analysis Program (Actimetrics) to determine clock oscillation period, length amplitude and phase. Briefly, raw data following the first cycle from day 2 to day 5 were fitted to a linear baseline, and the baseline-subtracted data (polynomial number = 1) were fitted to a sine wave, from which period length and goodness of fit and damping constant were determined. For samples that showed persistent rhythms, goodness-of-fit of >80% was usually achieved.

### siRNA and shRNA transfection, lentiviral plasmid construction and infection

siRNAs were purchased from Integrated DNA Technologies. Transfection were conducted using Lipofectamine RNAi MAX reagent (Invitrogen). 48-72 hours post transfection, cells were collected for protein or used for Lumicycle analysis. Lentiviral vectors expressing SRF or MRTF-A/B shRNA were obtained from Open Biosystems, as previously described (32). Three siRNA or shRNAs for each target were tested for knockdown efficiency. To generate shRNA knockdown lines, recombinant lentiviruses were produced by transient transfection in 293T cells using the calcium-phosphate method. Infectious lentiviral particles were harvested at 48 hr post-transfection to infect Per2-Luc fibroblasts. Two days post-infection, cells were selected with 2 μg/ml puromycin to obtain stable expression lines used for luminometry analyses. Lentiviral transduction efficiency was tested using lentiviral expressing GFP construct for GFP expression efficiency at nearly 95%.

### Immunoblot Analysis

20-40 µg of total protein were resolved on SDS-PAGE gel and transferred to PVDF membrane for immunoblotting (32). Immunoblots were developed by chemiluminescence kit (Pierce Biotechnology). Source and dilution information of primary antibodies are included as Suppl. Table S1. Appropriate specific secondary antibodies were used at a dilution of 1:3000.

### Quantitative real-time PCR analysis

RNeasy miniprep kits (Qiagen) were used to isolate total RNA from cells. cDNA was generated using q-Script cDNA kit (Quanta Biosciences) and quantitative PCR was performed on an ABI Light Cycler with SYBR Green (Quanta Biosciences). Relative mRNA expression was determined using the comparative Ct method to normalize target genes 36B4 as internal controls. Primers are designed using Primer Bank experimentally validated sequences. Primer sequences are listed in supplementary Table S2.

### Phalloidin Immunofluorescence Staining

Fluorescence staining of actin was performed using Alexa Fluor 488-conjugateed phalloidin, similarly as described (32). Briefly, cells were fixed with 3.7% formaldehyde and permeabilized using 0.1% Triton X-100. Alexa Fluor 488 phalloidin (10 µg/mL) at 1:200 dilution together with DAPI (1:2000) was incubated for 30 min at room temperature. Fluorescence images were taken using an ECHO microscope.

### ChIP-qPCR

Immunoprecipitation was performed using SRF or MKL1 antibody or control rabbit IgG with Magnetic Protein A/G beads (Magna ChIP A/G kit, Millipore), as described previously (36). Mouse myofibroblast chromatin was sonicated and purified following formaldehyde fixation. Real-time PCR was carried out in triplicates using purified chromatin with specific primers for predicted Bmal1 binding E or E’-box elements identified within the gene regulatory regions. Primers flanking known Bmal1 E-box in Rev-erbα promoter was used as positive control and TBP first exon primers as negative control. Fold enrichment were expressed normalized to 1% of input over IgG. ChIP primer sequences are listed in Supplementary Table S3.

### Statistical analysis

Data was expressed as mean ± SE. Differences between groups were examined for statistical significance using unpaired two-tailed Student’s t-test or ANOVA for multiple group comparison as indicated using Prism by GraphPad. P<0.05 was considered statistically significant.

## Acknowledgements

We thank Drs. Steve Kay and Meng Qu at the University of Southern California for sharing the luciferase reporter cell lines used in this study. KM is a faculty member supported by the NCI-designated Comprehensive Cancer Center at the City of Hope National Cancer Center. This project was supported by a grant from National Institute of Health 1R01DK112794 to KM. The funder had no role in study design, data collection and analysis, decision to publish, or preparation of the manuscript.

## Authorship Statement

XX and WL: data curation and investigation, formal analysis, manuscript editing; JN: data curation and manuscript editing; KM: formal analysis, project administration, manuscript writing and editing, and funding acquisition.

